# Cortical Changes Associated with Isha Kriya Meditation Revealed by Encephalography in Novice and Experienced Meditators – a Longitudinal Pilot Study

**DOI:** 10.1101/2024.12.30.630798

**Authors:** Ramana V. Vishnubhotla, Preeti U. Reed, Balachundhar Subramaniam

## Abstract

**Background:** Isha Kriya (IK) is a widely available meditation practice that incorporates breathing regulation that has shown to improve self-reported symptoms of stress, anxiety, and depression. An increasing amount of research has been published on the effects of various meditative practices on scalp electroencephalography (EEG). However, the effects of IK on cortical activity have not been reported previously.

**Methods:** Healthy volunteers aged 18 years or older were invited to participate. Participants were categorized as novice or experienced in meditation. EEG spectral features, computed during the eyes-closed condition before and soon after each IK meditation practice, were evaluated both at the start and after 6 weeks of IK meditation training.

**Results:** This longitudinal study examined the effects of IK meditation on cortical state and trait patterns in a cohort of eight participants who practiced IK meditation over a period of 6 weeks. Across the two sessions, a simultaneous increase in global periodic alpha power was observed in multiple subjects (N=6) but this was not observed in all subjects/sessions. We observed an increase in periodic theta band power, particularly in the frontal regions, which emerged as a common state effect in all participants. Longitudinally, we observed an increased periodic gamma power in the resting state EEG in all the experienced meditators in the parietal occipital region. The changes in novices on the other hand was in the alpha and beta bands.

**Conclusion:** Overall, in this pilot study, we report the changes in the quantitative EEG of the practitioners of Isha Kriya meditation over a 6-week cycle and investigated the difference between the start and end of that period at an individual level. We recommend future studies with a larger sample size and over a longer duration.

**Trial registration:** This trial is registered with the US National Institutes of Health on ClinicalTrials.gov with the trial identification number NCT03459690 on February 14, 2018.

## Introduction

Meditation as a practice and technique for training the mind has gained popularity as a research domain over recent years, in parallel with the growing traction it has gained within western society. This increase in interest is attributed to the various physical and mental benefits that meditation has shown to have on those who practice (1, 2). As a result of psychological, cognitive, and motivational traits, this form of mental training has impacted a practitioner’s perception, attention, emotional processing, and neuroplasticity, among many other components (2-4).

Two of the most commonly studied meditation practices include focused attention (FA) and open monitoring (OM) (2, 5, 6). FA involves maintaining selective or specific attention on an object or concept of choice, thereby anchoring their attention and awareness to focus. OM differs in that there is no specific focus on an object, rather attention to awareness itself, which allows for thoughts and sensations to flow through but also dissolve (7, 8). While FA represents a more acute focus, and OM a broadened focus, studies have observed that it is a combination of the two practices that contribute to mindfulness (3, 9). These practices have been applied in various clinical interventions, spanning from anxiety and depression, to chronic pain, and cancer, with multiple treatments, developed and programs being developed as a result (1, 10).

Studies have highlighted different aspects of how meditation impacts the brain and neural networking and have utilized neuroimaging techniques such as electroencephalography (EEG) and functional magnetic resonance imagining (fMRI) to further detail findings. EEG is non-invasive and provides a measure of neural network synchronization by analyzing spatiotemporal brain activity (5, 11). A common finding among many meditation studies has been the impact that meditation has on EEG alpha and theta activity and synchronization, demonstrating attention processing and an effect on specific regions and structures of the brain (12).

Different meditation traditions have been compared to one another, to determine how brain activity differs between the practices and experiences. In doing so, EEG readings have been interpreted beyond high and low frequencies, as seen in a 2017 study that looked specifically at the effects that particular practices have on brain activity and found that they all showed increased gamma amplitude. In addition to this positive correlation between gamma power and meditation experience, the increase was localized to the parieto-occipital region (13).

Neuroplasticity in meditation has also been seen with regards to the default mode network (DMN) and shown to have its particular regions deactivate or decrease in connectivity during mindfulness processes (2, 7). This observation has been noted in multiple studies and indicates that though the exact changes in the structure vary between practices, a correlation exists between the DMN and gamma oscillations, in relation to self-referential processing (4, 14).

In contrast to comparing various practices, studies have used EEG data to analyze how varying experiences within the same practice impact brain activity. Between the state of rest and a meditative state, different power increases have been observed, especially in regard to theta, alpha, and gamma bands. The differences in how the power increases have been shown to be dependent upon technique and experience with one meditation practice, and therefore also depict a relationship to neural processing that is specific to that practice (5, 10, 12). This was noted in a 2018 study by Kakumanu et al (15), in which they found that the type of meditative state of an experienced practitioner correlated with the changes in certain EEG powers and helped to determine what roles the state had in neural processing. Cognitively intense states due to sustained attention depicted an increase in low-alpha power, more effective states had greater high-theta power, and increased delta power was in association with enhanced internal processing. Power spectral changes in this study were linked to proficiency in practice rather than the duration of the technique.

While EEG serves to objectively measure the quality of various practices of meditation and mindfulness, no literature has been found examining the effects of Isha Kriya meditation on EEG brain waves. Isha Kriya is a practice that incorporates breathing regulation to produce a positive effect on the mind and body and does not require any complex training for practitioners. It has shown to decrease short-term stress (16) and reduce symptoms of anxiety and depression after weeks of practice (17). The goal of this study was to understand if practitioners of Isha Kriya meditation will show quantitative EEG changes over a 6-week cycle and investigate the difference between the start and end of that time period. Additionally, this study aimed to observe differences in EEG activity between experienced and novice Isha Kriya meditators. We recorded EEG measures in both novice and experienced meditators, once at baseline and once after 6-weeks of daily meditation and completion of a meditation diary. With the usage of online-guided tools for meditations, we hypothesized that the EEG readings of a novice practitioner after 6 weeks would reflect those of experienced practitioners, and further demonstrate the positive correlation between meditation and brain activity.

## Methods

### Study registration

This study was conducted in accordance with the Declaration of Helsinki (18). The study protocol was approved by the Committee on Clinical Investigations Institutional Review Board (IRB) at Beth Israel Deaconess Medical Center, Boston (Protocol number 2018-P-000040). This trial is registered with the US National Institutes of Health on ClinicalTrials.gov with the Trial identification number NCT03459690 on February 14, 2018. Healthy volunteers aged 18 years or older were invited to participate. The novice meditators (N) were defined as no meditation practice in previous years, objectively defined as <20 hours of lifetime meditation. On the contrary, an experienced meditator (E) was defined by >30 minutes of meditation practice for at least 5 days a week over the past year. In total, in this study data from nine volunteers (4 E and 5 N) participated in the study of which 2 participants (1 N, 1 E) did not participate in the recording session after 6 weeks and 1 participant’s recording during the first session stopped in between.

### Experimental protocol

The study participants were grouped into either novice meditators or expert meditators based on their meditation experience. Isha Kriya, a simple and easy to practice mindfulness technique taught by Isha Foundation, was introduced to both groups. Its simplicity and wide availability makes Isha Kriya a good choice for introducing meditation to beginners. EEG recordings were collected at baseline and then repeated at 6 weeks (+/-1 week). All enrolled participants were asked to practice Isha kriya meditation twice daily, for the study duration. Five minutes of eyes-closed resting-state data were obtained before (RS I) and after (RS II) the meditation. During the IK practice, participants either sat in a cross-legged posture or on a chair with their spine comfortably erect. Stage 1 involved focus on the breath while mentally focusing on two thoughts. Stage 2 involved uttering the sound followed by stage 3 where subjects sat with their eyes closed. The stages of the experimental protocol are illustrated in Figure 1. It is important to note that all participants were properly trained in the meditation before practicing. The ENOBIO EEG device (Neuroelectrics, Cambridge, MA) acquired 32 channel EEG recordings from the participants and the data was sampled at a rate of 500 Hz.

**Figure 1.**
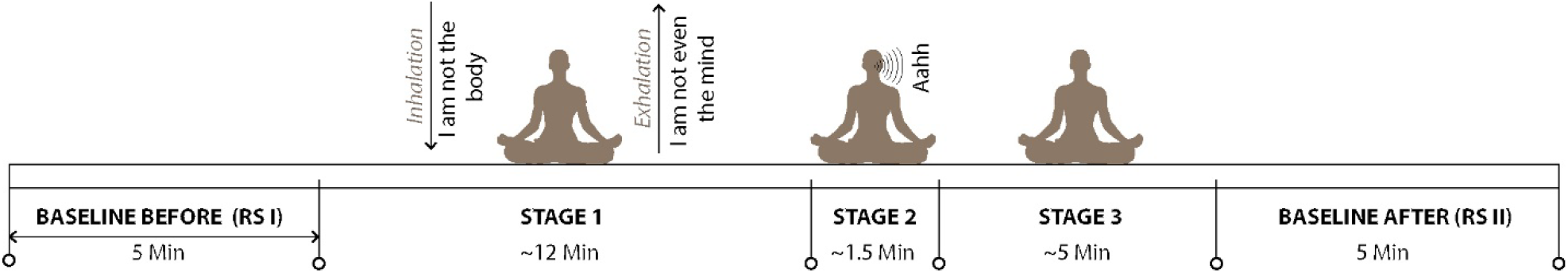
Experimental protocol

### Pre-processing

The preprocessing was implemented in MATLAB R2019b (MathWorks, MA) using custom functions and the EEGLAB toolbox (19). The preprocessing pipeline is summarized in Figure 2. The raw EEG was first high pass filtered using a 4th order zero-phase Butterworth filter with a cutoff frequency of 1 Hz followed by a low pass filter (4th order zero-phase Butterworth filter) with a cut off frequency of 95 Hz and an IIR notch filter at 60 Hz. The filters were implemented using the filtfilt function in Matlab to avoid phase distortion. The resting-state data (RS I and RS II) before and after the practice was initially segmented for further processing (Fig. 1). The data was then downsampled to 200 Hz.

**Figure 2.**
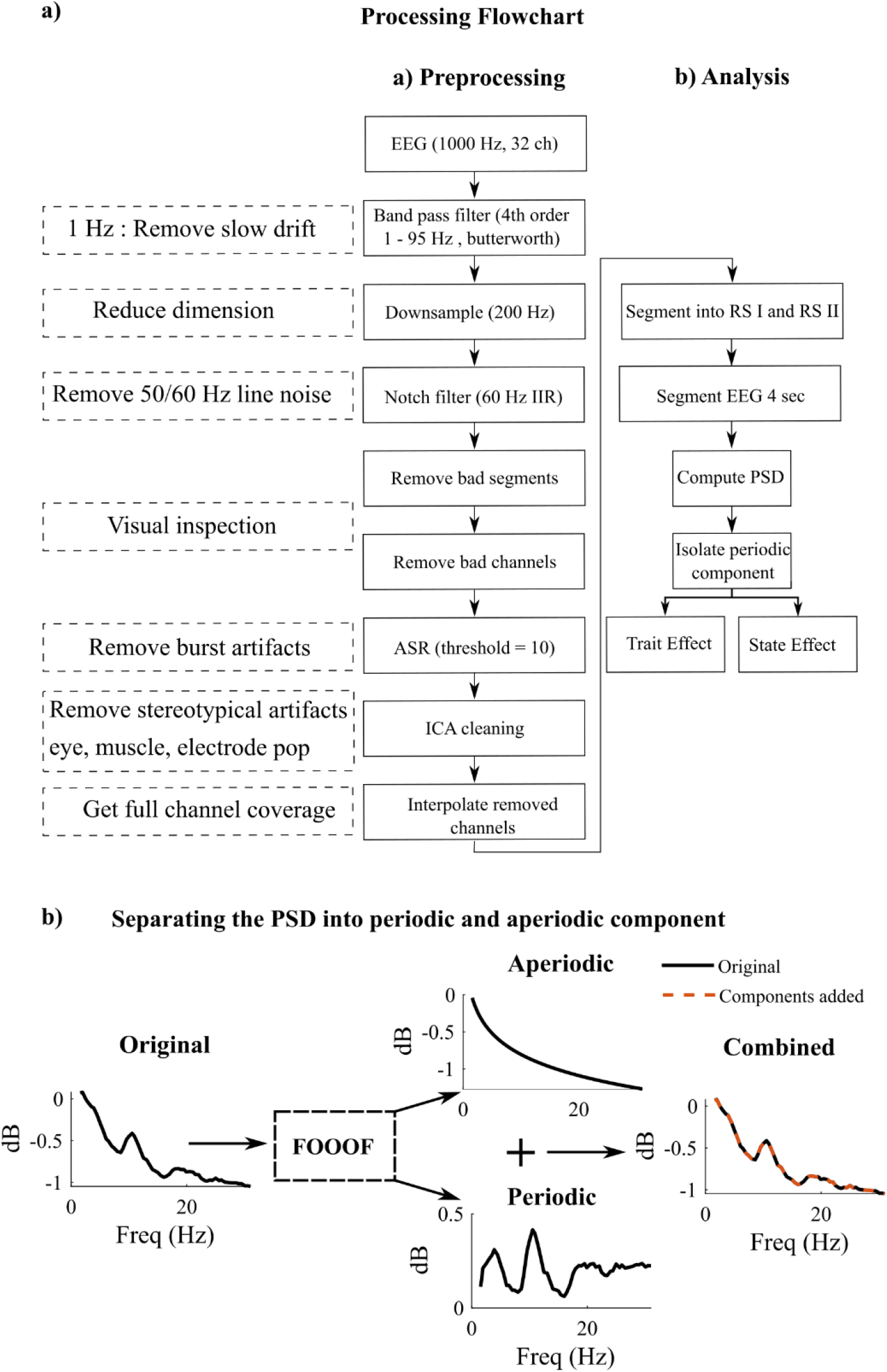
Processing flowchart. a) Cleaning and analysis pipeline followed in the paper on the EEG data segmented into RS I and RS II conditions b) FOOOF method applied on the PSD from one of the meditators (PO3 and PO4 mean).

Next, bad channels and windows were manually inspected and removed from all the subjects. Artifact subspace reconstruction was then performed to remove any large sudden bursts (20). A sliding window (length of 500 ms) and a threshold of 10 was used to identify corrupted subspaces for reconstruction. The thresholds were chosen based on empirical testing of multiple thresholds as well as based on recommendations from prior studies comparing the effects of these parameters (21). The signals were then decomposed into their independent components using Infomax independent component analysis (ICA) (22). This was done to remove eye, muscle, or any channel pops retained. Topoplot distribution, time-series data, and power spectral density were evaluated to identify these artifactual components and were later removed (discarded ICs: min 2; max 10).

### Spectral decomposition

Initially, the RS I and RS II periods were extracted to evaluate the state effects of the practice. These sections were then further segmented into 4-second windows without any overlap. The frequency power spectra in these windows in the frequency range from 1 Hz to 95 Hz with 0.5 Hz resolution was estimated using discrete prolate spheroidal sequences and the multi-tapers were computed in Fieldtrip (23). From the spectra, the absolute median band powers in the delta (1:4 Hz), theta (4:8 Hz), alpha (8:13 Hz), beta (13:30), gamma low (30:55 Hz) were computed. The relative power was then computed by normalizing each band power w.r.t. the total power in the window.

### Isolating periodic components

To better isolate the spectral peak components, the power spectral densities (PSD) were separated into periodic and aperiodic (1/f background noise) components using the FOOOF toolbox in python (Haller et al. 2018). It considers PSD to be a combination of aperiodic (*L*) and N oscillations above this aperiodic component referred to as peaks, modeled individually as a gaussian function *Gn*.

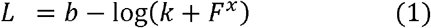

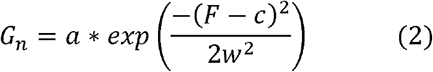

Here b is the broadband offset, X is the slope and k is the knee parameter and F is the vector of input frequencies. In Equation (2) a is the amplitude, c is the center frequency, w the bandwidth of gaussian. Aperiodic and periodic components in the frequency range of 1 to 55 Hz were estimated and the power in theta and alpha band in the periodic component was computed to evaluate the state effect as before.

### State effect: spectral change as a result of meditation

First, the state effects of IK on the spectral content in EEG, or how the power distribution in the different bands changed as a result of meditation, was assessed. Since there was a difference of approximately 20 minutes between the two baseline conditions, this could lead to a shift in total power not necessarily due to meditation. The time difference could cause spurious differences due to impedance change in either signal and reference electrodes or due to other environmental factors. To partially account for this, we did not use absolute power nor relative power. The supplementary section provides a case study on how information might be distorted when reporting these metrics using simulated power spectral densities. The section also shows how periodic power extraction could minimize these issues. The difference in the median power in the periodic component of the PSD between the RS II and the RS I is therefore computed. The changes will be reported for both pre and post 6-week training.

### Trait effect: longitudinal spectral changes in resting-state after 6 weeks

To evaluate whether the resting state EEG changes if a person does meditation for a continuous period of time, the periodic power was compared during the resting state eyes closed condition (RS1) for each of the 6 subjects who participated in the follow-up recording after 6 weeks. This quantifies how the power distribution during resting state changes as a result of 6 weeks of meditation practice. Considering the same reason as above, the analysis is limited to using the periodic power of the power spectral density.

### Statistical Testing

Subject level statistical testing here is done to test whether the power in windows post-meditation is significantly different from that in the baseline before the meditation. The normality of the data was evaluated using the one-sample Kolmogorov-Smirnov test. Since the test rejected the hypothesis that the data came from a normal distribution, non-parametric testing was performed by bootstrapping the band powers using permutations of independent t-tests with alpha = 0.05 in Fieldtrip (23). The *p* value is approximated using a Monte-Carlo estimate. A total of 2000 resample of the data were used to evaluate the significance similar to Braboszcz et al. (13). Due to the smaller number of subjects per group, the group level analysis is not performed in the study and effects will be assessed at an individual level.

## Results

### State effect

The change in mean periodic band power due to meditation is shown in Figure 3. In the subject level analysis, the power in the theta band increased in all participants, particularly after 6 weeks. The channels which had a significant difference in power at p < 0.05 between the two conditions are marked with circles in Figure 3. After 6 weeks, NS1 who did not have an increased theta power at the start of the training also exhibited an increase in theta power. On the other hand, a simultaneous increase in alpha power is observed in multiple subjects but was not common in all participants. It was seen in two out of three experienced meditators after 6 weeks. ES2 who had only elevated theta power at the start of the study showed a simultaneous increase in both theta and alpha after 6 weeks.

**Figure 3.**
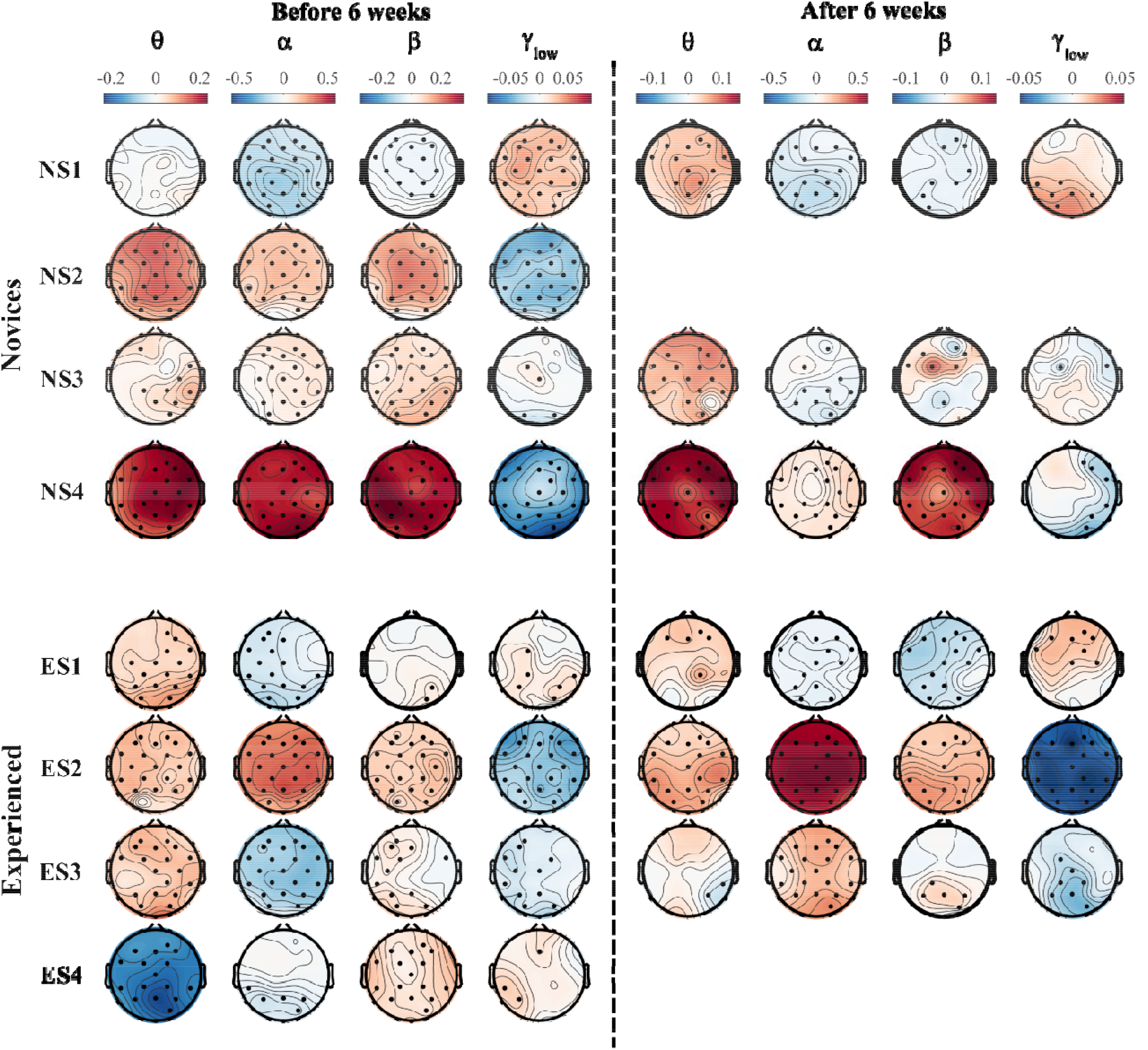
State effect: Change in periodic power due to meditation assessed both before and after 6 weeks; dots indicate the channels with a significant difference at p < 0.05.

Between the two sessions, three other novices also exhibited simultaneous alpha power increases. Beta power also increased in multiple subjects however gamma power generally was reduced during the meditation. Overall, the theta band power increase was the most consistent trend. Before 6 weeks, the theta power increased in six out of the eight participants whereas it reduced in two. The increase in the six subjects was statistically significant at p < 0.05. After the 6 weeks of practice, theta power increased in all the subjects of which five subjects had a statistically significant increase at p < 0.05. This increase was mainly localized in the frontal region.

### Trait effect

The change in periodic power in six of the participants who returned after 6 weeks is shown in Fig 4. In the subject level analysis, all the three experienced meditators, exhibited a significant increase in periodic gamma-band power in the resting state after 6 weeks, compared to the windows at the start of the study. The increase was mainly localized in the parietal occipital region. However, in the novices, the common change in power was largely in the alpha and beta band and not in the gamma band. The power in the alpha and beta frequencies in the experienced meditators reduced after 6 weeks.

**Figure 4.**
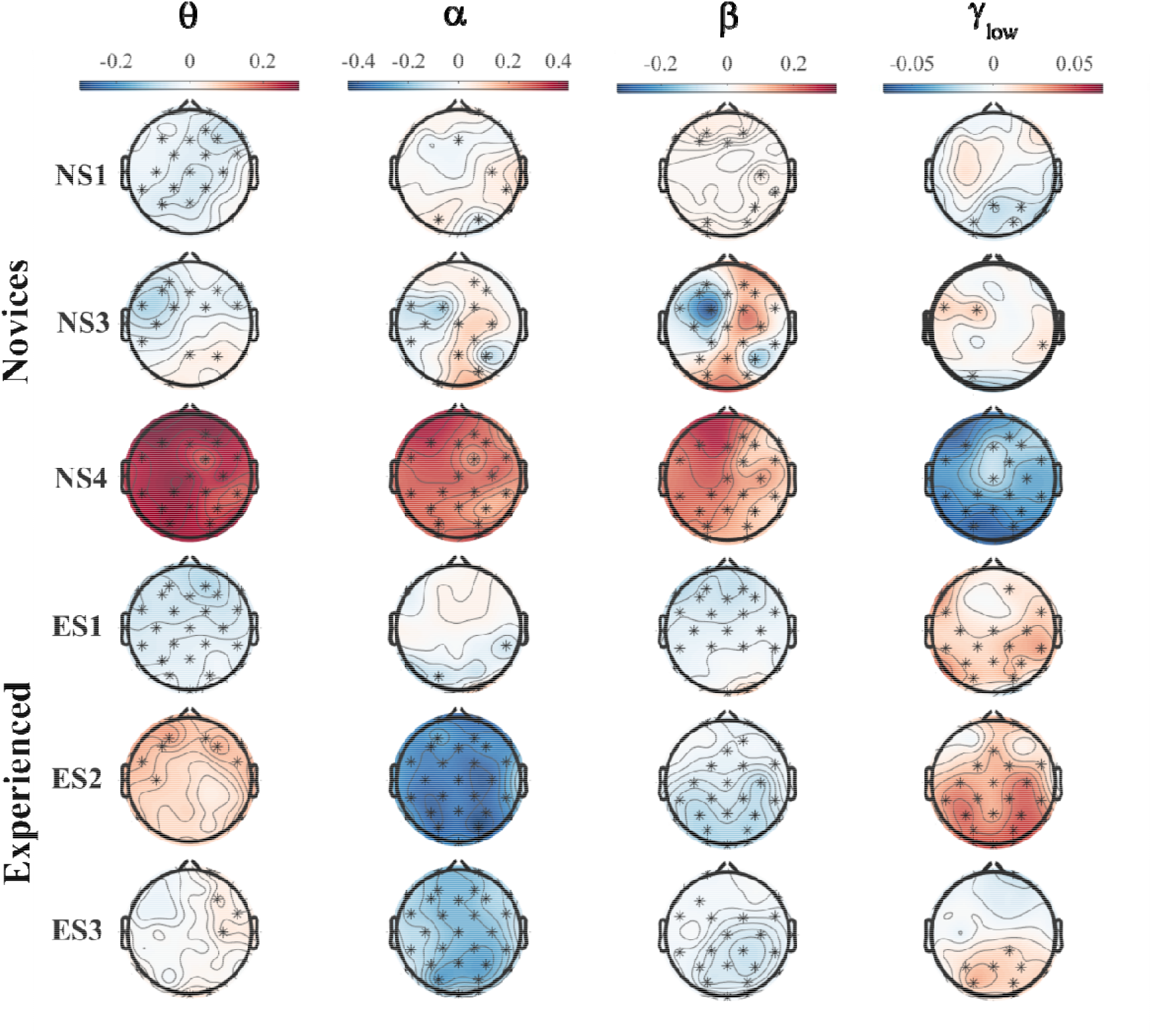
Trait effect. Longitudinal change in resting-state periodic power (during RS 1 period) after continuous IK practice for 6 weeks; * indicates the channels with a significant difference at p < 0.05.

### Discussion

This preliminary study observed the longitudinal changes from 6 weeks of practice of Isha Kriya in experienced and novice meditators. While both groups demonstrated some significant changes, the changes observed within each group was different. In the experienced meditator group, there was significantly greater gamma-band power at rest after 6 weeks. On the other hand, novice meditators demonstrated changes in alpha and beta power. The authors were expecting that EEG signal in novice meditators would be similar experienced meditators after six weeks, so these results were unexpected. The differences in changes in EEG signal based on meditator experience is noteworthy.

Even though cortical changes associated with meditation have been of interest for a long time, there exists no clear consensus on what is considered to be a generic trait and state effects (24). Among the changes reported, the modulation of theta and alpha power has shown some consistency across studies (5, 24). In our study as well, we observed the increase in periodic theta power to be the consistent trend in all the participants, particularly after 6 weeks of practice. The functional role of theta power in multisensory attentional processing and cognition has been studied extensively (25-27). Meditation too is a technique that heavily involves the attentional networks in the brain and an increase in theta power could be associated with this. Supporting this, prior studies have shown an overall increase in cerebral blood flow and activation in the anterior cingulate, dorsolateral prefrontal cortex and parietal regions during meditation, which are cortical structures heavily involved in the attention mechanisms (28, 29).

Though not all participants had an increased alpha activity, many of them did have a significant increase in both theta and alpha band power. An increase in alpha power is often associated with inhibitory actions by blocking task-irrelevant regions (30, 31). Therefore, the overall increase in periodic alpha coupled with the increased theta power might be associated with retaining a state of sustained attention by blocking external unwanted stimuli or information. The simultaneous increase in alpha was seen in two out of 3 experienced meditators after 6 weeks. The third experienced meditator (ES1), on the other hand, had a significant increase in power in the frontal gamma power with a decrease in alpha band power. Novice subject 4 also had a small increase in frontal gamma even though not significant during the second visit. In the second visit, NS4 had a lower increase in alpha band power relative to the first visit during which there was no increased gamma-band power. This could be indicative that the participant might be consciously thinking/distracted during that particular day causing a reduction in alpha power.

The state changes observed were quite variable across participants when evaluated using relative power as well as absolute power (supplementary materials). Prior investigations on the effects of meditation have reported similar variability and incongruencies in state effects in multiple studies (5, 32). One of the contributing factors for this could be the longer time separation between the two baseline conditions. This could lead to a change in absolute voltage values either due to a change in impedance at the reference &/ground electrode or the individual channels due to various environmental factors. Even though relative power partially accounts for this as itself normalizes to the power in the window, it might fail to capture simultaneous changes in multiple bands particularly if one band has significantly higher power. Here, we demonstrated (supplementary material) that separating the periodic component and evaluating the power after isolating the aperiodic component had a much stable and consistent state effect across the participants.

A long-term practice of meditation should induce changes in the resting state EEG patterns over time. Prior studies have reported the gamma-band power in the parietal occipital regions to be significantly higher in long-term meditators from 3 separate meditation traditions compared to the age-matched meditation naïve control group (13). The power also correlated with the hours of meditation, suggesting a potential trait effect. Thus, we expected after 6 weeks, the gamma power in all participants should increase. In this study, we did observe the resting state power in the gamma frequency band to increase in all experienced meditators after the 6 weeks. However, this was not observed in novice meditators in which power increase was mainly in the alpha/beta band and not the gamma band. The review paper by Cahn et al. has shown that studies that tracked longitudinal changes in EEG in novices, demonstrated an increase in alpha power at the end of 4-6 weeks. This suggests that there could be stages to how the resting state spectral features are modified as a result of long-term meditation practice. It could be that during the initial few months when someone is introduced to meditation, the change is reflected mainly in the alpha band, and it only translates to higher frequency bands after sufficient proficiency. Since the sample size and the tracked duration in the study is low, we cannot make this claim in this study. It would require looking at a larger cohort and tracking them for a considerably longer period to evaluate this hypothesis.

This being a pilot study, the study does suffer from multiple limitations. The number of participants present was low because of which the strength of the statistical test will be low and there is a possibility of false-negative observations. For the same reasons, we have not accounted for multiple comparisons and hence the results lack statistical strength. The results can therefore only be discussed qualitatively and future studies with a higher number of participants should be carried out to replicate the findings. In this study, the experienced meditators were meditation practitioners from other meditative traditions, and this would have confounded with how they perceived the IK practice. Future studies therefore should target recruiting participants who are experienced in IK practice alone to better evaluate stable state effects.

## Conclusion

Overall, this study performed a longitudinal EEG recording to assess the cortical changes associated with Isha Kriya; a popular yet simple meditation practice whose EEG spectral changes have not been studied in the past. This meditation program is available online for free which could lead to easier accessibility for a wider community. We identified that the commonly reported state effects of other meditation practices were shared by Isha kriya practice as well. The study identified the trend towards an increase in power in the upper gamma band in experienced meditators whereas the change was in the alpha and beta band for novices after the 6 weeks of practice. Future studies need to have larger sample sizes and should assess long-term changes in novice practitioners and determine if these changes are consistent with those observed in experienced meditators.

## Ethics Statement

This study was conducted in accordance with the Declaration of Helsinki (18). The study protocol was approved by the Committee on Clinical Investigations Institutional Review Board (IRB) at Beth Israel Deaconess Medical Center, Boston (Protocol number 2018-P-000040). All subjects provided verbal consent for the study.

